# Causal inference for the effect of environmental chemicals on chronic kidney disease

**DOI:** 10.1101/769430

**Authors:** Jing Zhao, Paige Hinton, Qin Ma

## Abstract

There is evidence from a limited number of statistical and animal studies that suggest that perfluoroalkyl acids (PFAs) are linked to a decline in kidney function. Thus, PFA exposure may be a modifiable risk factor for chronic kidney disease (CKD). As PFA is pervasive throughout our environment, determining its health effects is an important public health concern. We examined cross-sectional data from the 2009-2010 cycle of NHANES using generalized propensity score (GPS) analysis and univariate and multivariate ordinary least squares (OLS) regression to determine the link between urinary PFA concentration and estimated glomerular filtration rate (eGFR). GPS estimation methods used were Hirano-Imbens, additive spline, and a generalized additive model. Each of the statistical models used associated an increase in PFA concentration with a decline in eGFR, though the eGFR fit using the multivariate regression model were consistently higher than from the other four models. We conclude that PFA is a modifiable risk factor for CKD and GPS analysis produces credible results in estimating the effect of chemical exposures on continuous measure of kidney functions such as eGFR.

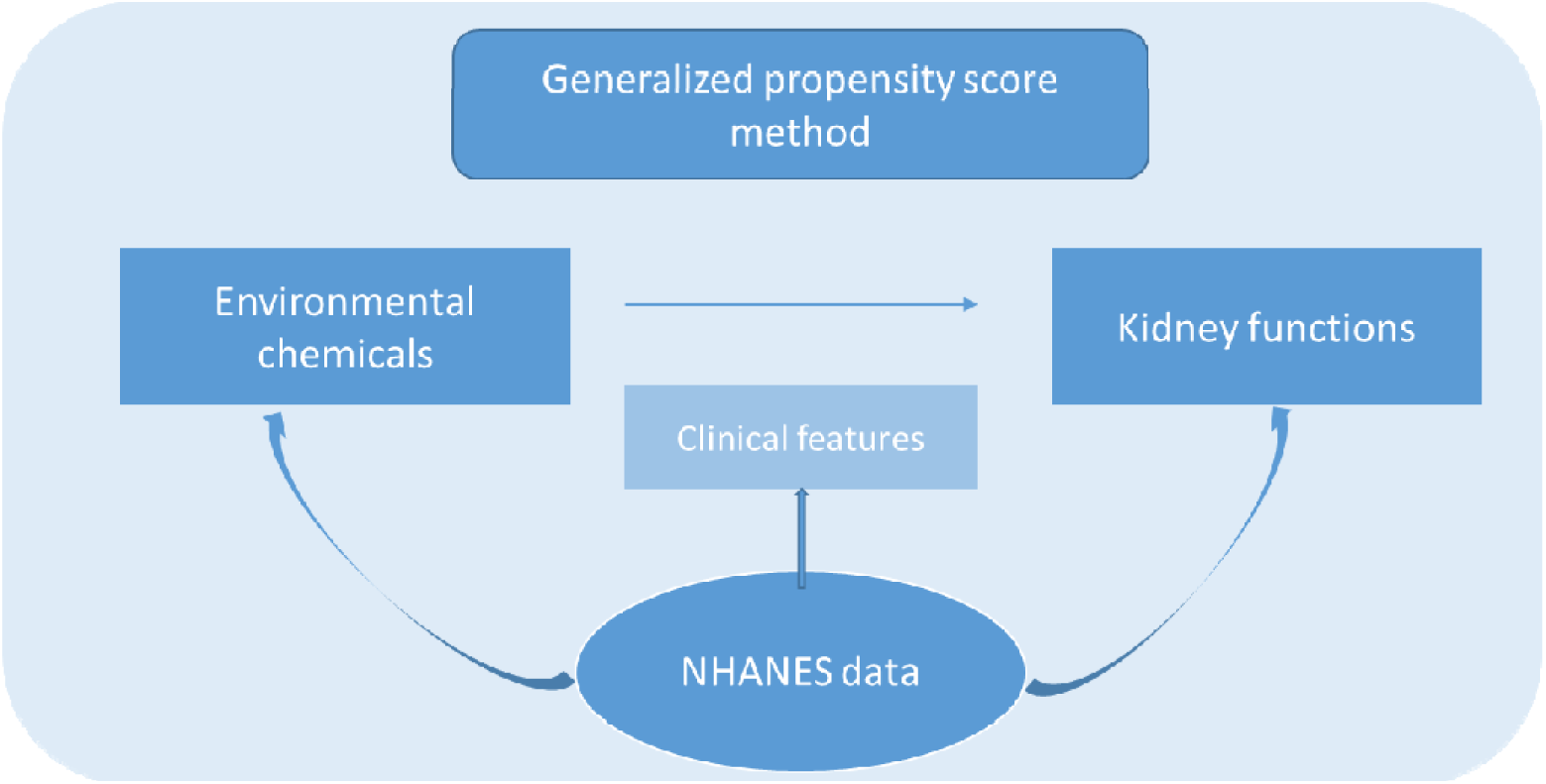
Graphical abstract.

## 1. INTRODUCTION

The prevalence of Chronic Kidney Disease (CKD), defined by an estimated glomerular filtration rate < 60 mL/minute/m^2^, continues to increase, and is present in about 13 % of the adult US population [1]. The presence of CKD raises the risk of other diseases including cardiovascular disease and kidney failure [1]. It is an early stage of renal disease, so it is possible to prevent or delay the progression of CKD to end-stage renal disease (ESRD) if detected early enough. Early intervention in kidney disease would alleviate patient suffering and the economic burden of treating the disease incurred by kidney transplant operations and dialysis required to treat ESRD.

Perfluoroalkyl acids (PFAs) are synthetic chemicals that have been detected in the blood of most people in the United States[2]. Animal studies suggest that there may be an association between PFAs and CKD, but human research studies on the topic are scarce. PFA levels in the blood have been positively associated with CKD but support from more evidences needed [2]. PFAs are of concern because they are persistent environmental contaminants that bioaccumulate in organisms and are biomagnified along food chains [3]. PFAs are found in a variety of products, including food packaging, surfactants, lubricants, sealants, stain-resistant sprays for textiles, and fire-retarding foams. Two PFAs, perfluorooctanoic acid (PFOA) and perfluorooctane sulfonate (PFOS), bioaccumulate in the kidneys, as kidneys are the primary route of elimination of PFAs. PFAs are associated with changes in the permeability of endothelial cells, which is considered a mechanism of renal failure based on rat models[4]. PFOA and PFOS have been previously associated with increased cholesterol levels, insulin resistance and risk of metabolic syndrome, which have been linked independently to an increased risk of CKD [2]. Increased serum levels of PFOA linked to decreased renal function (a decreased eGFR) and increased risk of having CKD, but reverse causality cannot be ruled out. Elevated serum PFA levels may be caused by decreased kidney function, instead of PFA causing a decline in kidney function[5]. PFAs have also been associated with hyperuricemia, insulin resistance, diabetes risk, and metabolic syndrome, each of which is an additional risk factor for CKD [5].

To intervene, we need to identify those individuals most at risk for the disease. An individual’s genetic and phenotypic characteristics both affect their risk in developing kidney disease, including genetic mutations, a family history, gender, ethnicity, age, obesity, socioeconomic status, smoking, nephrotoxins, acute kidney injury, diabetes mellitus, and hypertension [6]. Low birth weight is another risk factor for CKD because restricted intrauterine growth is linked with a low nephron number, which may lead to hypertension and kidney disease later in life due to the decreased GFR and increased albumin-to-creatinine ratio [6]. Obesity is one of the top risk factors for CKD, as increased glomerulus size and hyperfiltration increases pressure on the capillary wall, increasing the risk of kidney injuries. Kidney damage may be caused by complications of obesity, including inflammation, oxidative stress, endothelial dysfunction, and hypervolemia [6]. People with a history of acute kidney injury (AKI) are ten times more likely to develop ERSD within a year of their AKI diagnosis than those without an AKI [7]. Diabetes mellitus is another primary risk factor since kidney disease arises by hyperfiltration injury, oxidation, and advanced glycosylation end products [8]. Systemic hypertension leads to intraglomerular capillary pressure, which leads to, glomerulosclerosis, the hardening of the kidney, which eventually leads to kidney failure [8].

The risk of kidney disease varies among different demographic populations. ESRD occurs more frequently in men, but CKD occurs more frequently in women [6]. The discrepancy may be due to the confounding by other lifestyle factors [9]. There is an increased risk of ERSD in African Americans; one of the reasons may be that a mutation in the APOL1 gene associated with a significant risk in ERSD is only found in those of African descent [6]. Other social and environmental factors, along with possible genetic predispositions, are also likely contributors to the differences in risk of developing ESRD between African-American and non-African-American populations [10]. Kidney function decreases as age increases, and thus, elderly people are more likely to develop CKD [6]. Socioeconomic status is inversely associated with CKD risk. For example, in one study unemployed non-Hispanic blacks and Mexican-Americans were found to be twice as likely to have CKD than if they were employed [11]. However, opposing evidence exists that there was no statistical significance between a family history of ERSD and socioeconomic status after accounting for patients with a family history of ESRD at the community level [12], and more research needs to be done.

Lifestyle behaviors also affect the development of kidney disease. Smoking indirectly increases the risk of CKD by causing oxidative stress, endothelial dysfunction, glomerulosclerosis, and tubular atrophy [13]. Consumption of nephrotoxins, such as alcohol, recreational drugs, heavy metals, and analgesic drugs in excess, are associated with CKD [14]. The negative impact of environmental chemicals on human health has drawn much attention, with liver and kidney being the two major organs harmed by environmental chemicals [4].

Many of the risk factors that likely contribute to kidney disease are associated with one another, and it is difficult to determine if kidney disease is a direct or indirect consequence of a specific risk factor. For example, socioeconomic status does not directly cause chronic kidney disease, but factors associated with lower socioeconomic status, like lack of health care, poor dietary choices, substance abuse, and behavioral patterns increase the risk of developing and/or dying from CKD [10]. Addressing the ancestral causes of risk factors may have a greater positive effect on the treatment and outcome of patients with chronic kidney disease than each risk factor individually. In this study, we are aiming to elucidate the causal impact of environmental chemicals, PFA particularly, on kidney functions by controlling confounding factors using data from the National Health and Nutrition Examination Survey (NHANES). The data collection, data manipulation and statistical methods are demonstrated in Materials and Methods section. Data description and results for comparison between linear regression methods and generalized propensity score methods are showcased in the Results section. The achievements and limitations of this study is discussed in the last section.

## 2. MATERIAL AND METHODS

### 2.1 Data collection

We used the 2009-2010 cycle of NHANES [15] to study the statistical causal association between environmental chemicals and kidney function. Bisphenol A (BPA), polyaromatic hydrocarbons (PAHs), perfluoroalkyl acids (PFAs), phthalates, dioxins, furans, and polychlorinated biphenyls (PCBs) were the chemicals considered. The amount of each chemical in each subject was measured using the sum of the urinary concentration of each chemical’s respective metabolites. The metabolites of BPA, PAHs, PFAs, and phthalates were recorded as individual level data, but dioxins, furans, and PCBs were in the pooled format, and were not analyzed. We pulled out the environmental chemical data that were measured on the individual level (not pooled). We converted the concentration of each urinary metabolite to nanomoles per liter (nmol/L). The total concentration of PAH was calculated as the sum of the urinary concentrations of 2-hydroxyfluorene, 3-hydroxyfluorene, 9-hydroxyfluorene, 1-hydroxyphenanthrene, 2-hydroxyphenanthrene, 3-hydroxyphenanthrene, 1-hydroxpyrene, 1-naphthol, 2-naphthol, and 4-hydroxyphenanthrene. The total concentration of PFAs was calculated as the sum of the concentrations of perfluorooctanoic acid, perfluorooctane sulfonic acid, perfluorohexane sulfonic acid, 2-(N-Ethyl-perfluorooctane sulfonamido) acetic acid, 2-(N-Methyl-perfluorooctane sulfonamido) acetic acid, pefluorodecanoic acid, perfluorobutane sulfonic acid, perfluoroheptanoic acid, perfluorononanoic acid, perfluorooctane sulfonamide, perfluoroundecanoic acid, and perflurododecanoic acid. The total concentration of phthalates was split into high-molecular weight (HMW) and low-molecular weight (LMW) phthalates. The concentration of HMW phthalates was taken as the sum of the concentrations of mono(carboxynonyl) phthalate, mono(carboxyoctyl) phthalate, mono-2-ethyl-5-carboxypentyl phthalate, mono-(2-ethyl-5-hydroxyhexyl), mono-(2-ethyl)-hexyl phthalate, mono-(2-ethyl-5-oxohexyl), and mono-benzyl phthalate. The concentration of LMW phthalates was taken as the sum of the concentrations of mono-*n*-butyl phthalate, mono-(3-carboxypropyl) phthalate, monoethyl phthalate, mono-*n*-methyl phthalate, and mono-isobutyl phthalate.

Kidney function is measured by the glomerular filtration rate (GFR) clinically as a comprehensive index. Instead of directly being measured, GFR is estimated from serum creatinine and adjusted by age, gender and race [16]. Thus, in this study, kidney function of participants were represented by eGFR values, which were estimated using the CKD-EPI creatinine equation (2009) [16]:

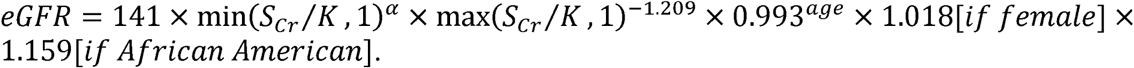

In order to establish the causal relationship between chemicals and kidney function, we extracted the confounding factors, including age, gender, race, systolic and diastolic blood pressure, income-to-poverty level ratio, BMI, diabetes status, smoking status and alcohol consumption habits. We excluded subjects with missing data for chemicals or confounding factors from our study.

### 2.2 Statistical methods

The concentrations of environmental chemicals including PFA, BPA, PAH, LMW phthalates, and HMW phthalates were log transformed to reduce the skewness of their distribution. The log-transformed chemical concentrations were analyzed as continuous variables. A univariate analysis was performed using t-test or chi-square test as appropriate to compare each of the characteristics from two groups defined by the first and fourth quartiles of the chemical level. Descriptive and univariate analyses were conducted using R Studio 3.4.4.

In this study we applied a generalized propensity score (GPS) method in the analysis of the effect of PFA on renal function and compared the performance of this method with linear regression models. The GPS is a continuation of the propensity score method and is the probability of an individual having a certain level of urinary PFA, given their baseline covariates. This removes bias introduced by the imbalance of baseline covariates regarding PFA concentrations, and differences in eGFR can be solely attributed to differences in PFA concentrations.

The GPS was first introduced in Hirano and Imbens 2004 [17]. The Hirano and Imbens method gives a parametric estimator that first estimates the GPS by regressing on the covariates and then estimates the eGFR based on the observed PFA concentration and fitted GPS [18]. The additive spline estimator estimates the ADRF in a two-step semiparametric process as outlined in Bia et al [19]. The first step is to parametrically model the GPS based on the covariates and create additive spline bases for the GPS and PFA. The eGFR is regressed on PFA, PFA bases, GPS, and GPS bases. The second step is to calculate the eGFR based on each PFA value and corresponding covariates, and then to average the eGFR to get the ADRF at that treatment value. The generalized additive model (GAM) estimator uses a treatment formula to estimate and model the GPS, and then the eGFR is estimated based on the treatment and the spline base terms from fitting the GPS [20].

The formulation of the Hirano and Imbens method in this study is outlined below:

For each unit *i* = 1, …, *N*, there exists a set of potential eGFR values, *Yi*(*c*), for some *c ∈ C*, where *c* is the treatment level. This gives the unit-level chemical response function. The average dose-response function (ADRF) is given by *µ* = *E*[*Y*_*i*_(*c*)]. Each unit *i* is associated with a vector of covariates, ***X***_***i***_, and the corresponding treatment level, *c*_*i*_ *∈* [*c*_0_, *c*_1_]. The variables *X*_*i*_, *c*_*i*_, and *Y*_*i*_ (*c*_*i*_) are observed.

The unconfoundedness assumption made by Rosenbaum and Rubin in 1983 is generalized for the GPS by Hirano and Imbens [17], which means if we condition on covariates, response to treatment level is independent of the treatment assignment, and bias due to varying covariate distributions for different PFA levels is reduced. This allows us to make comparisons in the eGFR between subjects with similar covariate values at different PFA levels.

As has been defined in [18], the conditional density function of the treatment given the covariates is *r*(*c, x*) = *f*_*C*|*X*_ (*c, x*) and the generalized propensity score is *R* = *r*(*C, X*).

The GPS has a balancing property, and the probability of treatment level does not depend on ***X***. Combining the balancing property with the unconfoundedness assumption, the level of PFA is unconfounded based on the GPS.

The GPS model was implemented and the average dose-response function (ADRF) was estimated using the *causaldrf* package. A variety of estimators for the ADRF in the *causaldrf* package were tested, including Hirano and Imbens (H-I), generalized additive model (GAM), and additive spline. The covariates adjusted for in the estimation of the ADRF were the diabetes status, systolic blood pressure (SBP), diastolic blood pressure (DBP), ratio of family income-to-poverty level (socioecon), body mass index (BMI), number of weekly drinks, and smoker status. These estimators were compared to two OLS regression models. One OLS model regressed eGFR on the log PFA concentration, diabetes status, SBP, DBP, socio-economic status, smoker status, the interaction between log PFA and SBP, and the interaction between log PFA and socioeconomic status. The other OLS model was a simple univariate regression of eGFR on log PFA concentration.

## 3. RESULTS

### 3.1 Comparison between linear regression methods and GPS methods

In observational studies, there are two main methods in elucidating the associations between predictors and outcomes: non-causal methods such as linear regression and causal methods such as propensity score analysis. Non-causal methods, e.g. multivariate regression analysis has a couple of advantages over propensity score analysis. One is that it can detect an interaction effect between the treatment variable and other covariates. Another is that the effect of a confounder on the outcome variable can also be estimated. Propensity score analysis, on the other hand, is only able to estimate an average effect of treatment on outcome [21]. If there are many pre-treatment variables that are potential confounders, conventional methods of covariate adjustment might be insufficient. Propensity scores can theoretically remove all bias associated with differences in pre-treatment variable distributions across different treatment levels [22]. It has the logical advantage of separating subjects based on their propensity score, grouping them based on similar covariate values, and comparing outcomes between subjects within the same propensity score group with varying exposure levels. Within each group of propensity scores of similar values, any difference in outcome can be attributed to differences in exposure level. From an analysis point of view, it is necessary in both propensity score analysis and linear regression that the model be correctly specified to deduce any significant associations between exposure and outcome, but it is easier to run diagnostics on propensity score models than on regression models. In causal inference, it is important to ensure covariate overlap between individuals across exposure levels, whereas with regression, residuals need to be checked. Propensity score analysis is more objective than regression in the sense that it is not influenced by the exposure variable. This objectivity is not possible in linear regression analysis.

We further discussed different literatures in revealing associations between environmental chemical - PFA exposure and renal outcome-eGFR using non-causal and causal inference models in Table 1. Shankar et. al used multivariate and age and sex adjusted regression models to investigate the relationship between serum PFA levels and eGFR and found that PFA and eGFR were negatively correlated with each other, even after adjusting for confounders [2]. As NHANES is population-based and representative in nature, confounders could be adjusted for. The data used was cross-sectional, preventing them from ruling out reverse causality. It is noted that in the case of reverse causality, the association between PFA and eGFR would still be significant from a public health standpoint [2]. In a different study, high serum levels of perfluorooctanoic acid (PFOA) and perfluorooctanesulfonic acid (PFOS), two PFAs, were associated with decreased eGFR in children and adolescents using univariate, bivariate, and weighted multivariate regression. This study also found differences in PFA concentration and eGFR by social and behavioral characteristics, emphasizing the need for confounding adjustment. The main limitation in the current literature examining the effect of serum PFA on eGFR using linear regression is the cross-sectional nature of the data used. As noted previously, propensity score analysis is unable to detect effects of explanatory variables other than PFA concentration. These findings were also based on cross-sectional NHANES data, preventing reverse causality from being ruled out and highlighting the need for longitudinal studies on the effect of PFAs on renal function. Results may also have been hindered by the lack of availability of potential confounders, such as lifestyle choices and concentrations of other environmental chemicals in the serum [23]. A study by Watkins et al also found an inverse association between eGFR and serum PFA concentration in a cross-sectional study of children and adolescents but suggests that reverse causation may play a part. This study was also done using linear regression [5]. Propensity score analysis is not able to address the reverse causality problem associated with cross-sectional data, and assumes all relevant confounders are adjusted for.

**Table 1.** Comparison between non-causal and causal methods

|  | Non-causal | Causal |
| --- | --- | --- |
| Input/<br>metho<br>d | 1. Standard survival analysis;<br>2. baseline covariates;<br>3. regression models;<br>4. cross-sectional data;<br>5. stratification by eGFR [24]. | 1. Propensity score [17];<br>2. stratification, logit;<br>3. longitudinal data;<br>4. baseline covariates. |
| Output | Odds ratios [2]. | Hazard ratio [25]. |
| Advantages | 1. Can show association between predictor variables and an outcome variable;<br>2. confounding adjustment. | 1. Simulate randomized controlled trial by minimizing baseline confounding;<br>2. confounding adjustment. |
| Limitations | 1. Causation of effect cannot be determined;<br>2. cannot rule out reverse causation [4, 5]. | 1. Cross-sectional data such as NHANES may be inappropriate for causal inference, as assumptions about temporal sequence are made [26]. |

### 3.2 Baseline characteristics and univariate analysis: Inverse relationship between urinary PFA concentration and eGFR

Characteristics of the 1607 participants in NHANES with laboratory test results available on PFA are shown in Table 2. PFA concentrations were summaries over demographic features, clinical measurements as well as CKD. The first and fourth quartiles of PFA were also compared against these variables. As showcased in Table 2, there is a statistically significant decrease in the eGFR of patients of the fourth quartile of urine PFA concentration (mean = 84.88) compared to those in the first quartile (mean = 101.79). People with higher PFA concentration (who were in the fourth quartile of PFA concentration) tended to be older, male, non-Hispanic black, have a lower income-to-poverty level ratio, consume more alcoholic beverages, have diabetes, and have higher systolic blood pressure than those with lower PFA concentration (who were in the first quartile). Based on a significance level of 0.05, there is a statistically significant difference between eGFR for the first and fourth quartiles of urinary PFA concentrations. A decrease in renal function measured by eGFR was associated with an increase urinary PFA concentration. We also performed analyses on the association between urinary concentrations of bisphenol A, polyaromatic hydrocarbons, and phthalates and eGFR, but no statistically significant results were found. This may be due to unconsidered confounders or interactions with other chemicals. The summary statistics of all the other chemicals can be found in the supplementary file.

**Table 2:** Baseline characteristics and univariate analysis

| Perfluoroalkyl acids (PFA) | total, n = 1607 |  | 1st quartile, n = 403 |  | 4th quartile, n = 401 |  |  |
| --- | --- | --- | --- | --- | --- | --- | --- |
| Characteristic | mean (SE) | n (%) | mean (SE) |  | mean (SE) | n (%) | p-value |
| Age | 49.93 (0.44) |  | 43.24 (0.81) |  | 58.02 (0.85) |  | 2.20E-16 |
| Male sex |  | 756 (47.04) |  | 90 (22.33) |  | 255 (63.59) | 2.20E-16 |
| Race (non-Hispanic black) |  | 263 (16.37) |  | 60 (14.89) |  | 87 (21.70) | 0.016 |
| Ratio of family income to poverty level | 2.52 (0.04) |  | 2.13 (0.08) |  | 2.73 (0.08) |  | 7.68E-08 |
| Smoker |  | 364 (20.44) |  | 79 (19.60) |  | 67 (16.71) | 0.33 |
| Weekly alcoholic beverages | 1.03 (0.05) |  | 0.56 (0.06) |  | 1.49 (0.11) |  | 3.25E-12 |
| Body mass index | 29.24 (0.17) |  | 29.70 (0.40) |  | 28.83 (0.31) |  | 0.87 |
| Diabetes mellitus |  | 170 (10.58) |  | 28 (6.95) |  | 53 (13.22) | 4.60E-03 |
| Systolic blood pressure (mm Hg) | 122.85 (0.46) |  | 118.74 (0.93) |  | 128.21 (0.88) |  | 4.10E-13 |
| Diastolic blood pressure (mm Hg) | 69.24 (0.33) |  | 69.59 (0.69) |  | 68.64 (0.71) |  | 0.336 |
| serum creatinine (mg/dL) | 0.89<br>(0.01) |  | 0.83<br>(0.03) |  | 0.96<br>(0.01) |  | 1.05E-04 |
| estimated glomerular filtration rate (mL/minute/ 1.73 m<sup>2</sup>) | 93.02<br>(0.58) |  | 101.79<br>(1.21) |  | 84.88<br>(1.07) |  | 2.20E-16 |
| Chronic kidney disease |  | 139<br>(8.65) |  | 21<br>(5.21) |  | 60<br>(14.96) | 7.60E-06 |

### 3.3 Association between PFA concentration and eGFR using GPS and regression methods

The common support plot for the GPS methods is shown in Figure 1, which demonstrates that there are distinct overlaps in the range of propensity scores across different PFA concentration levels. The extent of overlap between each pair of level 1 vs. level 6, level 2 vs level 5, and level 3 vs. level 4 was satisfactory.

**Figure 1.**
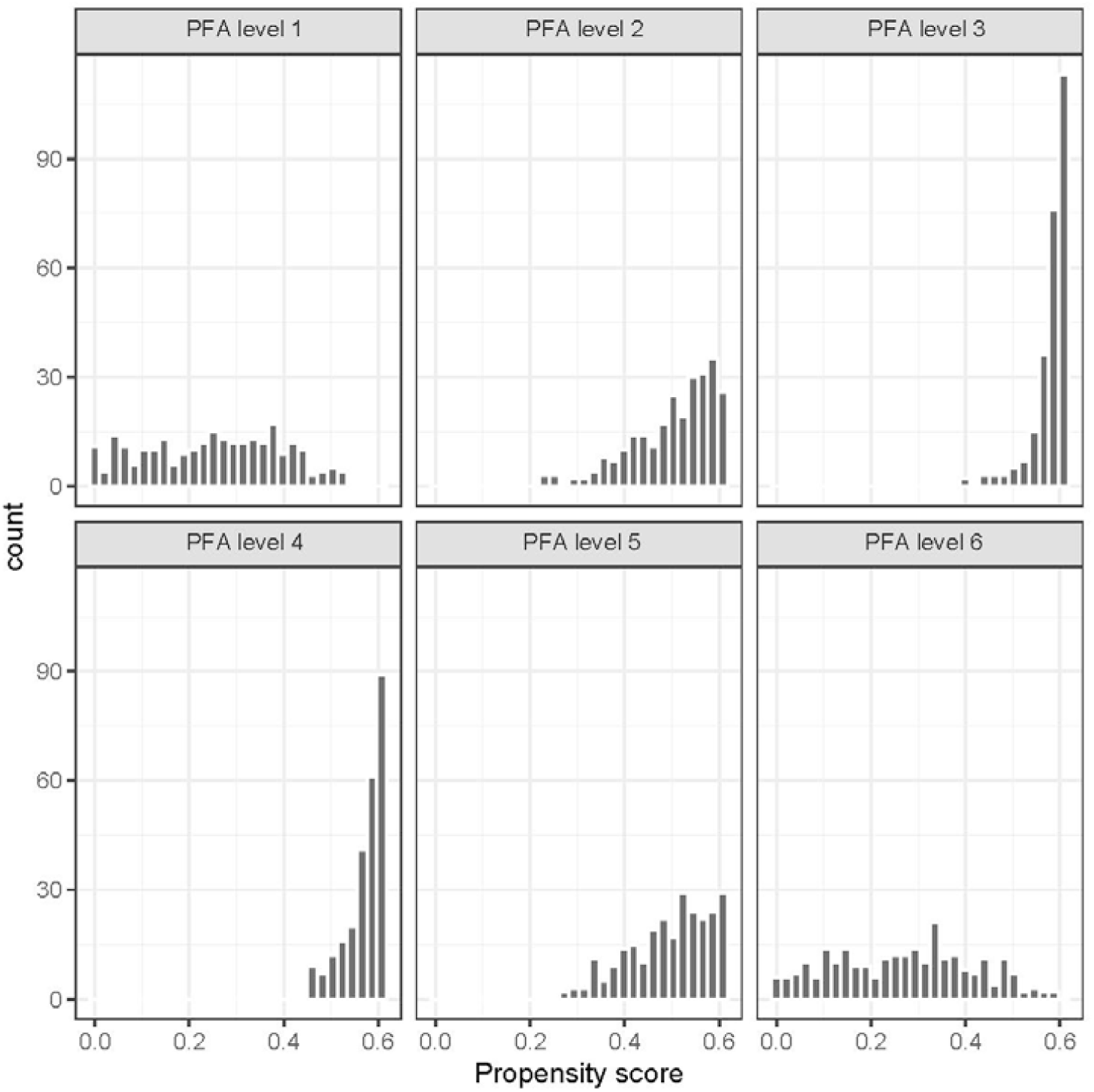
Distribution of Propensity Score across Treatment Groups: PFA levels are ordered as concentration ranges from low to high.

Both non-causal (regression) and causal methods (GPS) were applied in estimating the effect of PFA on eGFR. Table 3 contains diagnostic statistics for three different estimators of the dose-response function of PFA concentration of eGFR. The models based on a GPS and the regression models all show a significant relationship between eGFR and the regressor variables, including PFA. The R^2^ values for each of the five models are relatively low, with the univariate regression model having the lowest (R^2^ = 0.053) and the multivariate regression having the highest (R^2^ = 0.21).

**Table 3.**
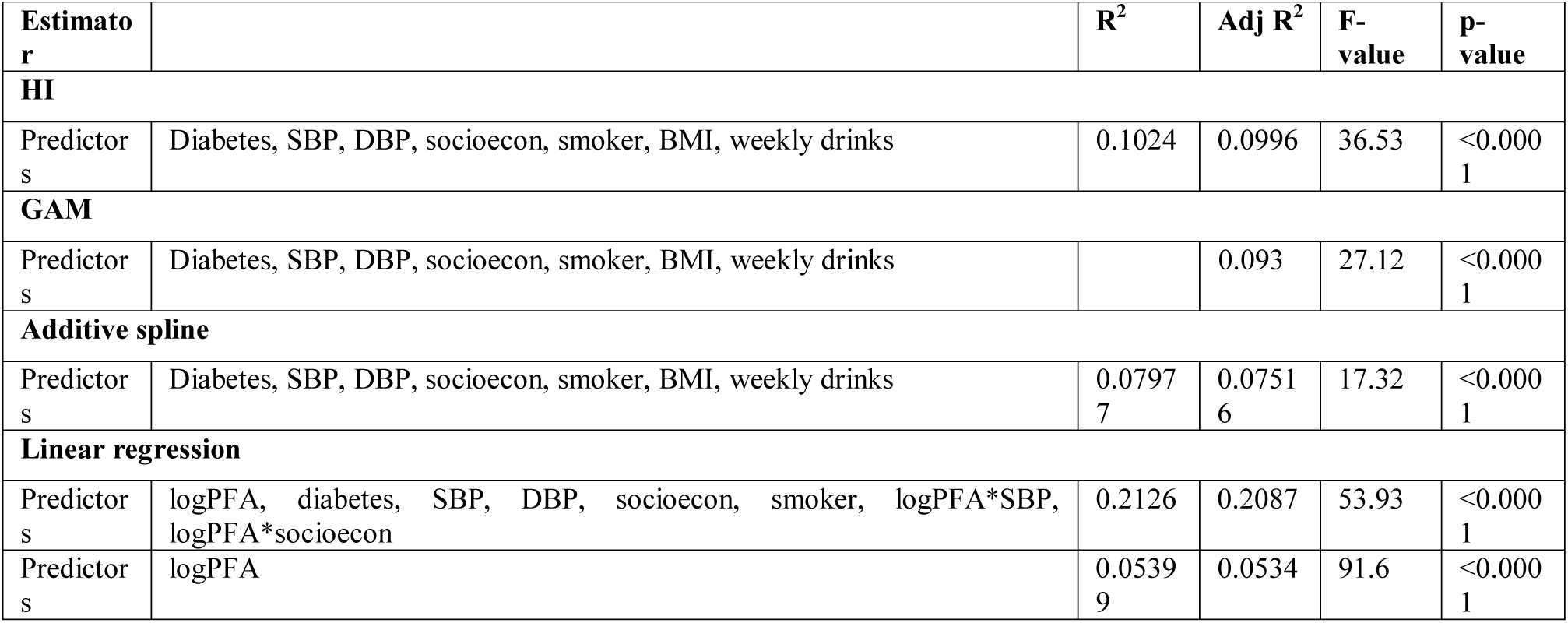
Diagnostic statistics of ADRF estimators and OLS regression.

| Estimator | | $R^2$ | Adj $R^2$ | F-value | p-value |
| --- | --- | --- | --- | --- | --- |
| **HI** |  |  |  |  |  |
| Predictors | Diabetes, SBP, DBP, socioecon, smoker, BMI, weekly drinks | 0.1024 | 0.0996 | 36.53 | <0.0001 |
| **GAM** |  |  |  |  |  |
| Predictors | Diabetes, SBP, DBP, socioecon, smoker, BMI, weekly drinks |  | 0.093 | 27.12 | <0.0001 |
| **Additive spline** |  |  |  |  |  |
| Predictors | Diabetes, SBP, DBP, socioecon, smoker, BMI, weekly drinks | 0.07977 | 0.07516 | 17.32 | <0.0001 |
| **Linear regression** |  |  |  |  |  |
| Predictors | logPFA, diabetes, SBP, DBP, socioecon, smoker, logPFA*SBP, logPFA*socioecon | 0.2126 | 0.2087 | 53.93 | <0.0001 |
| Predictors | logPFA | 0.05399 | 0.0534 | 91.6 | <0.0001 |

We further predicted eGFR based on the value of the log PFA for our five different estimators as shown in Figure 2: GAM, additive spline, and H-I, which employ the GPS, and univariate and multivariate regression. The three GPS models are qualitatively similar in shape, increasing with log PFA before reaching a peak, decreasing, and then beginning to plateau. The univariate OLS regression model shows similar results to the GPS models, but predicts lower eGFR at lower log PFA values and slighter higher eGFR as log PFA increases. The multivariate OLS model was plotted using the estimated coefficient for log PFA. The multivariate regression model estimates much higher eGFR values across log PFA values.

**Figure 2.**
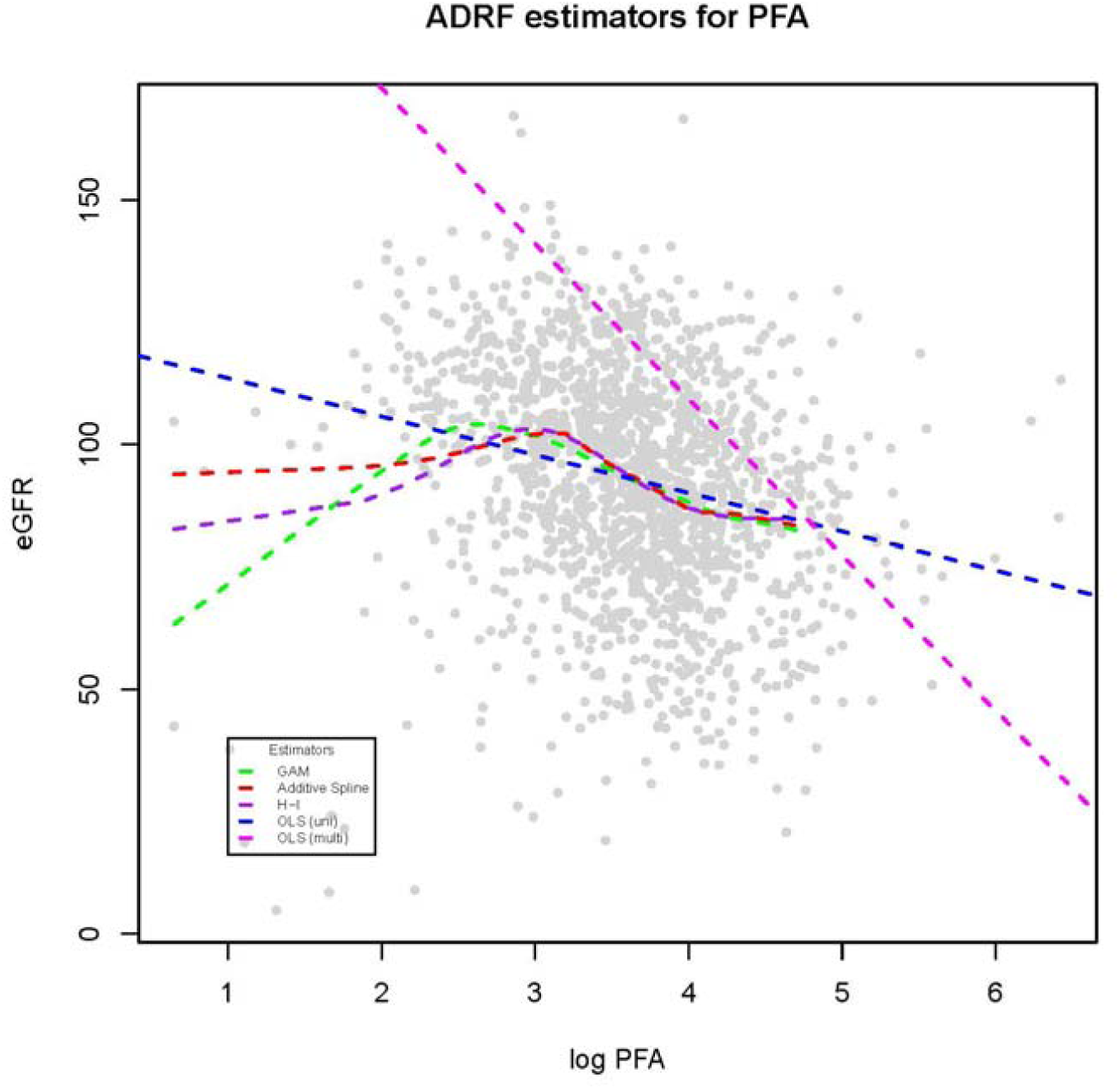
Estimated eGFR corresponding to log PFA concentration levels using GPS and regression.

## 4. DISCUSSION AND CONCLUSIONS

PFA is hypothesized to have a negative effect on renal function and supported by animal studies. But evidences from human studies are very rare in suggesting that PFA is linked to a decline in kidney function. In this study, observational data from the 2009-2010 cycle of NHANES were used in elucidating the statistical causal association between urinary PFA concentration and eGFR using GPS methods. Three GPS methods were examined and the fitted eGFR values over PFA concentration demonstrate an increase in PFA concentration with a decline in eGFR. Thus we conclude that PFA is a modifiable risk factor for CKD and GPS analysis produces credible results in estimating the effect of chemical exposures on continuous measure of kidney functions such as eGFR.

This study has significant impacts on determining the health effects of environmental chemicals. Environmental chemicals such as PFA are invading human life in an unobservable manner and experiments in evaluating the health effects of these chemicals are impractical to implement in clinical settings. Thus advanced statistical techniques such as GPS methods provide us with a way of causing no additional risks to patients while eliminating the biases introduced by unbalanced covariates in observational studies. Results from causal methods as GPS also facilitate biomedical researchers formulating more solid hypotheses in examining the health related environmental chemicals.

However, limitations still exist in the current study. One of the limitations was the lack of availability of data. We were unable to investigate potential interaction effects on kidney function between PFA and other chemicals, including BPA, PAH, and phthalates, as the subset of individuals in NHANES 2009-2010 with recorded PFA levels did not have levels measured for these other chemicals. Also, as NHANES data is cross-sectional in nature, we could not evaluate the effect of PFA on individuals over time, which would be ideal when studying a chronic disease such as CKD. In addition, although the GPS methods used in this study have satisfactory common support which means most of the participants’ data were used in the analysis, some of the data has been discarded due to no matched sample in the comparison group. To overcome these limitations, we are planning to use methods such as inverse probability weighting which can improve the efficiency and reduce bias of unweighted estimators. We will also reach out and request access to databases that have paired and longitudinal data sets.

We have shown that GPS-based methods can be used to identify a causal relationship between two continuous variables, and has theoretical advantages over OLS regression. Future work entails the investigation of the effect of interaction between PFA and other chemicals on CKD, and the application of causal inference methods to longitudinal studies, which would make an even stronger case for the negative effect of PFA on kidney function.

## Appendix

### 1. Literature review of BPA, PAH, phthalates, dioxins, furans, PCB, and glyphosate

Bisphenol A (BPA) is a synthetic estrogen used to in the production of polycarbonate plastics and epoxy resins, which are used in many consumer and industrial products [24]. Polycarbonate plastics are used in food packaging, safety equipment, and medical devices. Epoxy resins are used to coat metal surfaces, such as cans, water supply pipes, and bottle tops. BPA is a possible endocrine disruptor [27]. Dialysis patients have increased BPA exposure due to the use of BPA in dialysis tubing [4]. Increased levels of urinary BPA were associated with higher levels of albuminuria. Lower levels of urinary BPA excretion were associated with a decreased GFR, but the same study found no association between BPA excretion and BPA [24]. BPA is cleared rapidly by the kidneys [24]. Animal studies found an association between serum BPA and albuminuria and suggest that BPA negatively affects the glomerulus [4]. Few studies have looked exclusively at the relationship between BPA and estimated glomerular filtration rate (eGFR). One study found that a decrease in BPA and triclosan excretion was associated with a lowered GFR using data from NHANES 2003-2006 [24]. Elevated urinary BPA levels may increase the risk of hypertension and diabetes mellitus, two risk factors also associated with renal function [4].

Phthalates are separated into two categories based on molecular weight: low or high. Low molecular weight (LMW) phthalates are found in personal hygiene items such as shampoo, lotion, and cosmetics, while phthalates of high molecular weight (HMW) are involved in the production of vinyl plastics found in flooring, intravenous tubing, and food packaging [4]. Phthalates are possible endocrine disruptors, and infants and children may be more affected by phthalates because of the increased food consumed-to-body weight ratio. Diethyhexyl phthalate (DEHP) is a HMW phthalate used in medical supplies to administer blood and nutrition intravenously and respiratory gases to patients. Patients with chronic kidney disease (CKD) are at a high exposure risk to phthalates due to the number of medical treatments they undergo, such as dialysis, and the prevalence of phthalates in equipment used to complete such procedures. Albuminuria, the presence of albumin in the urine, is a symptom of kidney disease and a risk factor for cardiovascular disease. An increase in DEHP metabolites in the urine was associated with an increase in the albumin concentration in the urine, though LMW phthalates did not affect albuminuria [28]. An increase in DEHP metabolites in urine was associated with an increase in the systolic blood pressure, but phthalates of LMW had no effect [28].

Dibenzo-*p*-dioxins and dibenzo-furans are renal toxins that hyperuricemia (elevated levels of uric acid in the blood). Increased serum levels of furans are associated with high levels of serum uric acid levels. Hyperuricemia has previously been linked to diabetes mellitus and coronary artery disease, which are both risk factors for CKD[29]. There is an inverse relationship between polychlorinated dibenzo-p-dioxin (PCDD) and polychlorinated dibenzo-p-furan (PCDF) serum levels and eGFR. Dioxins and furans are toxic, persistent environmental chemicals arising from the production of pesticides and wood pulp that are biomagnified in the food chain[4]. The greatest source of dioxin exposure to humans is animal products, as dioxins accumulate in fat. Dioxins have been associated with cardiovascular disease, diabetes, and hypertension [4]. Exposure to PCDD/Fs may increase the risk of insulin resistance[29]^6^.

Polyaromatic hydrocarbons are produced from burning coal, oil, or gas, from grilling meat over charcoal, or in tobacco. Oxidized PAHs may be mutagenic. PAH exposure increases the risk of Balkan endemic nephropathy, a disease also associated with urothelial cancer [4]. PAHs also have a direct relationship with systolic and pulse pressure, but further studies are needed to confirm the hypertension risk associated with PAH exposure [4].

Polychlorinated biphenyls (PCBs) were used in the capacitors and coolants of electrical equipment until 1979 in the United States, when their persistence and ability to bioaccumulate in the tissues of organisms became apparent. Because of their environmental persistence, careless disposal practices, and use of products that contain them, PCBs are still affecting human populations today. No association was found between serum PCB levels and kidney function in a study done on sewage workers with elevated serum PCB levels who cleaned up sewage containing PCB in the municipal waste system. Further studies should focus on the effect of elevated PCB levels on kidney function and albuminuria. Elevated PCB serum concentration is also associated with hyperuricemia and hypertension, especially systolic blood pressure [4].

Glyphosate is a broadleaf herbicide used both on agricultural crops and on residential lawns. It now represents more than half of all active ingredients used in agricultural herbicides and is still growing in prevalence. As glyphosate-based herbicides (GBHs) are sprayed on a variety of crops, including maize, corn, wheat, and beans, residues of glyphosate and its primary metabolite AMPA have been found on harvested crops and even in processed foods. Part of the reason glyphosate residues are found on crops at harvest is the use of GBHs close to harvest time to desiccate crops and thus allow harvest to begin sooner. Glyphosate may be especially harmful to the kidneys and liver in comparison to other tissues. One study found there was a 10- to 100-fold increase in glyphosate levels in the kidneys and liver compared to fat, liver, and muscle [28]. Glyphosate has been associated with oxidative damage to the kidneys in rats [11]. Another study found an association between farmers who sprayed glyphosate, drank hard water, and developed CKD. The metal-chelating property possessed by glyphosate is the suspected explanation, but further investigation is needed to determine causation [13].

### 2. Baseline covariates comparisons of the first and fourth quartiles in BPA, PAH, phthalates, dioxins, furans, PCB, and glyphosate

Compared to those with urinary BPA concentrations in the first quartile, subjects with BPA concentrations in the fourth quartile were more likely to be younger, non-Hispanic black, have a lower ratio of family income-to-poverty level, and have a higher BMI. Subjects with higher urinary PAH levels were more likely to be younger, non-Hispanic black, have a lower income-to-poverty level ratio, smoke, and have a higher BMI than their counterparts with lower serum levels of PAH. Subjects with high HMW phthalate concentrations had a higher probability of being younger, male, have a higher BMI, and have a higher systolic blood pressure than those with low levels of serum HMW phthalates. Subjects with high LMW phthalates were more likely to be non-Hispanic black, have a lower income-to-poverty level ratio, and have a higher BMI. In the case of PAH, LMW phthalates, and HMW phthalates, eGFR values was higher in those with higher urinary chemical concentrations.

**Table 1:**
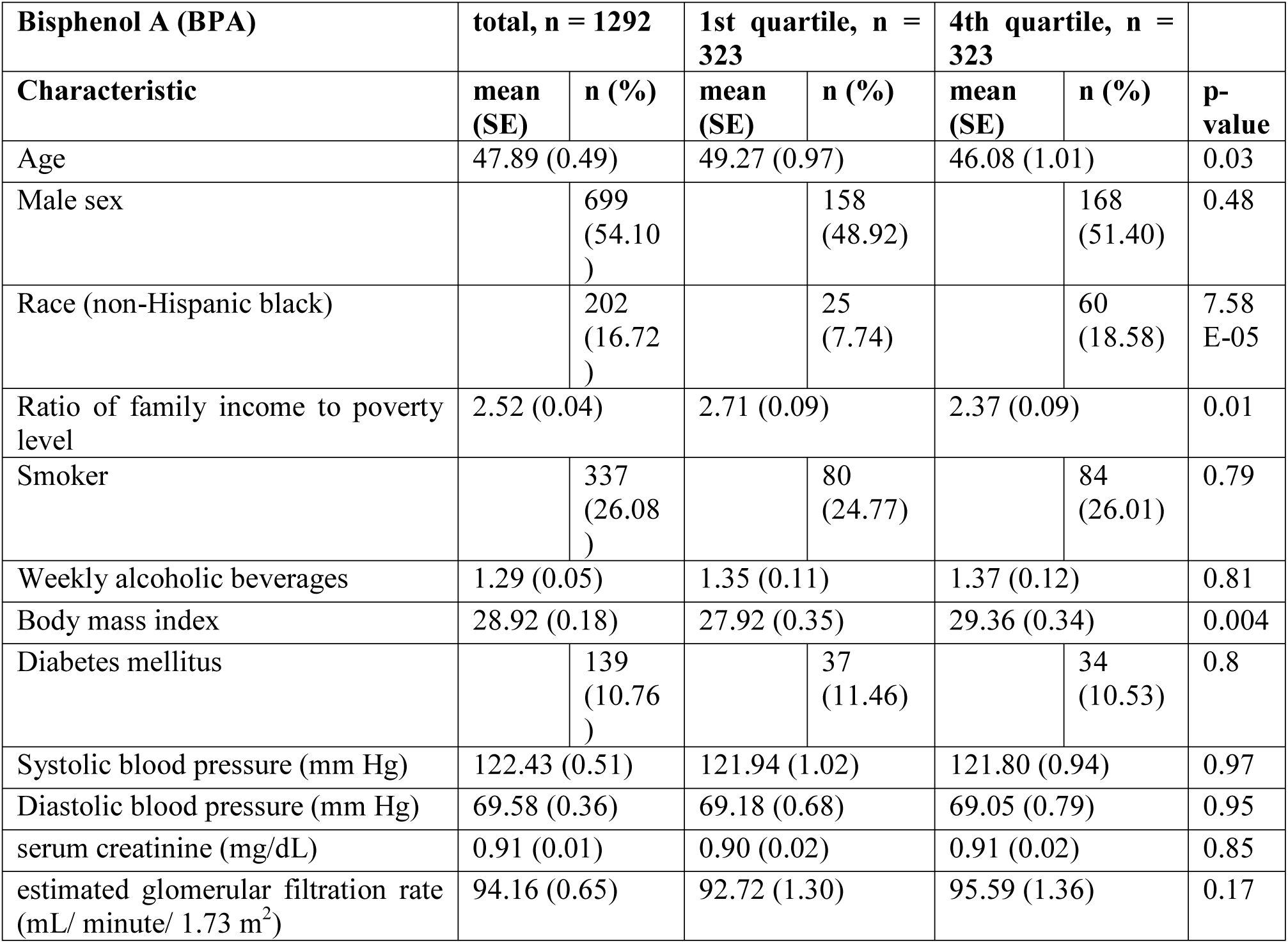
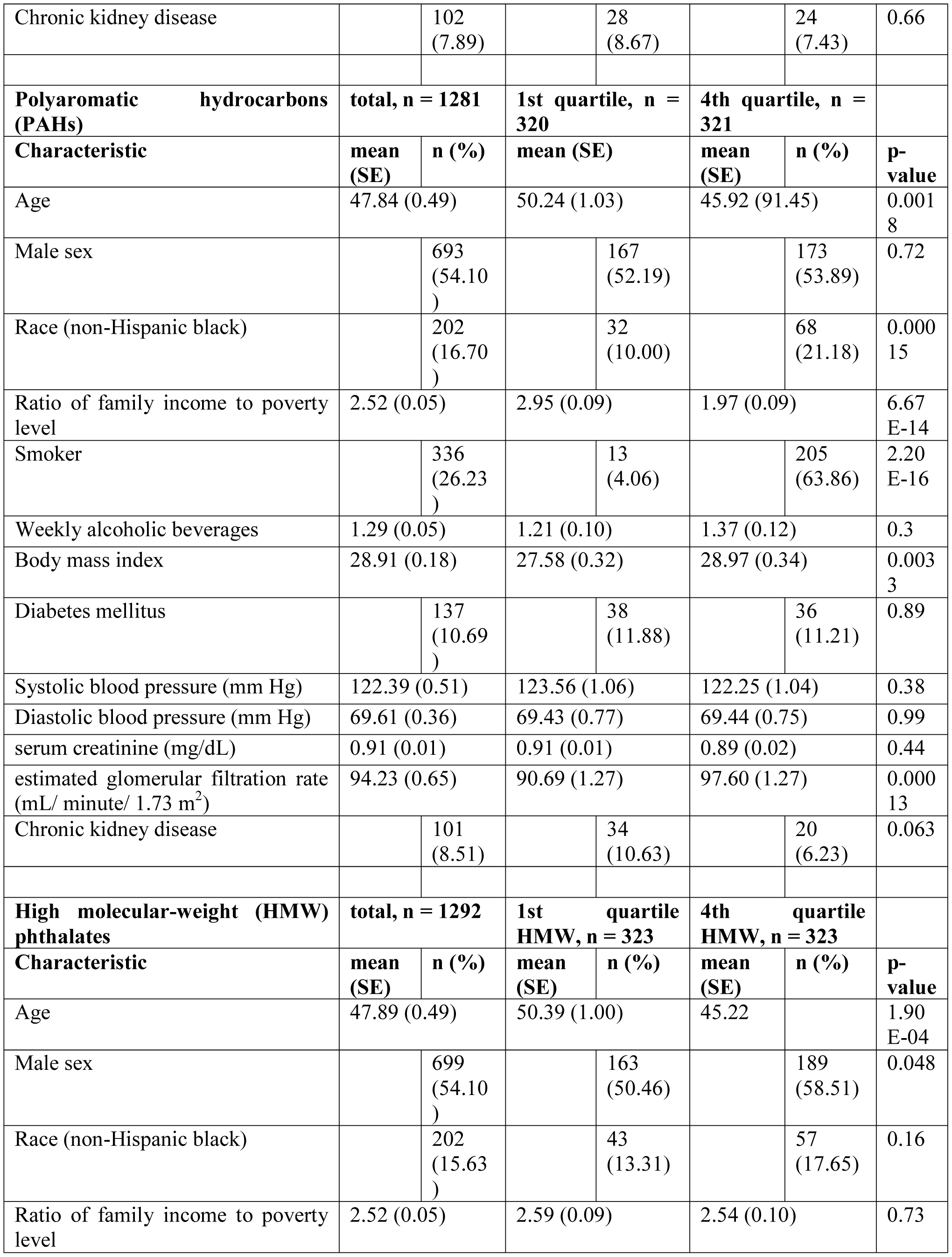
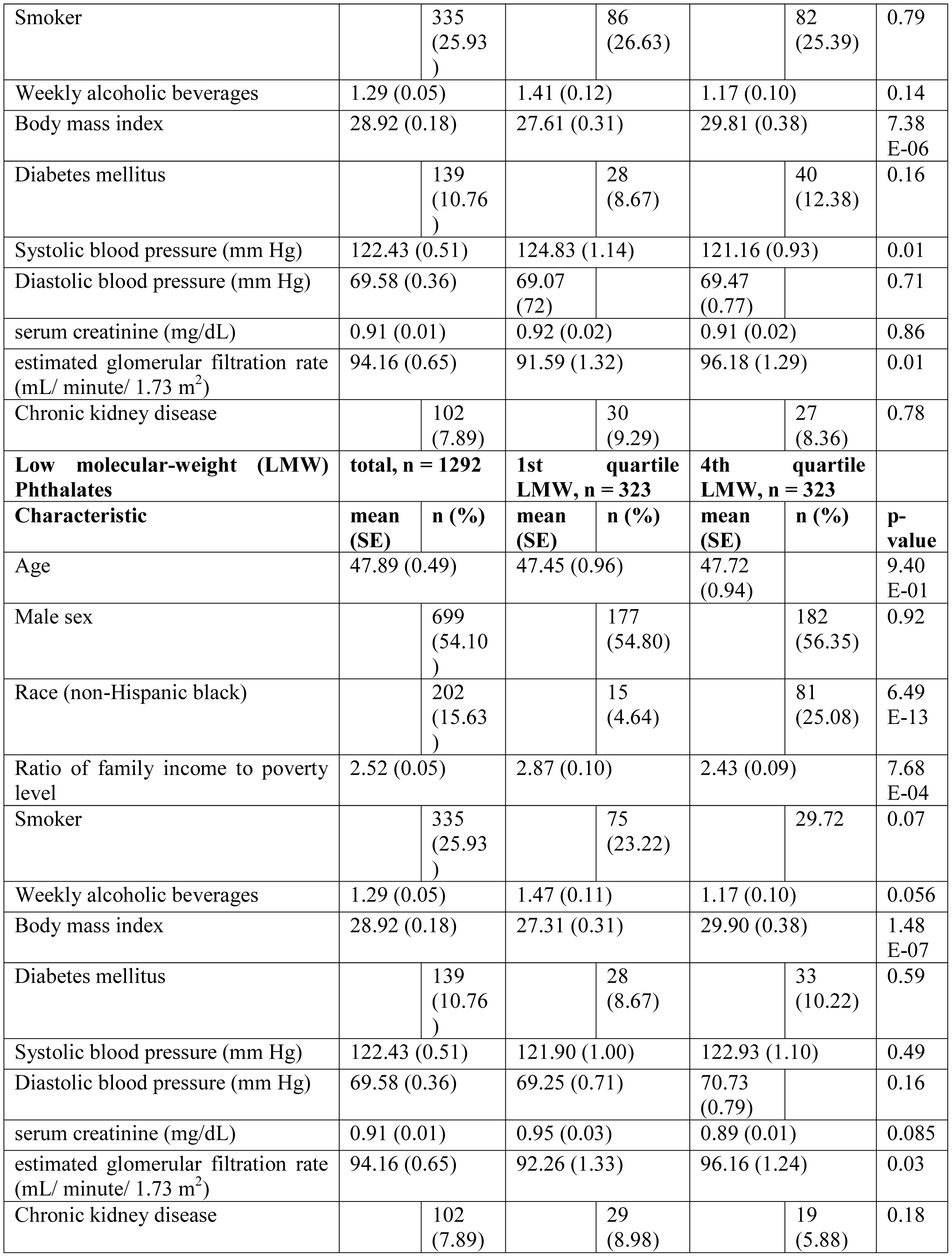
Covariate statistics in total and divided into quartiles with test results of comparing first and fourth quartiles

