## Appendix for "Causal inference for the effect of environmental chemicals on chronic kidney disease"

Table 1. Covariate statistics in total and divided into quartiles with test results of comparing first and fourth quartiles

| **Bisphenol A (BPA)** | **total, n = 1292** | | **1st quartile, n = 323** | | **4th quartile, n = 323** | |  |
| --- | --- | --- | --- | --- | --- | --- | --- |
| **Characteristic** | **mean (SE)** | **n (%)** | **mean (SE)** | **n (%)** | **mean (SE)** | **n (%)** | **p-value** |
| Age | 47.89 (0.49) | | 49.27 (0.97) | | 46.08 (1.01) | | 0.03 |
| Male sex |  | 699 (54.10) |  | 158 (48.92) |  | 168 (51.40) | 0.48 |
| Race (non-Hispanic black) |  | 202 (16.72) |  | 25 (7.74) |  | 60 (18.58) | 7.58E-05 |
| Ratio of family income to poverty level | 2.52 (0.04) | | 2.71 (0.09) | | 2.37 (0.09) | | 0.01 |
| Smoker |  | 337 (26.08) |  | 80 (24.77) |  | 84 (26.01) | 0.79 |
| Weekly alcoholic beverages | 1.29 (0.05) | | 1.35 (0.11) | | 1.37 (0.12) | | 0.81 |
| Body mass index | 28.92 (0.18) | | 27.92 (0.35) | | 29.36 (0.34) | | 0.004 |
| Diabetes mellitus |  | 139 (10.76) |  | 37 (11.46) |  | 34 (10.53) | 0.8 |
| Systolic blood pressure (mm Hg) | 122.43 (0.51) | | 121.94 (1.02) | | 121.80 (0.94) | | 0.97 |
| Diastolic blood pressure (mm Hg) | 69.58 (0.36) | | 69.18 (0.68) | | 69.05 (0.79) | | 0.95 |
| serum creatinine (mg/dL) | 0.91 (0.01) | | 0.90 (0.02) | | 0.91 (0.02) | | 0.85 |
| estimated glomerular filtration rate (mL/ minute/ 1.73 m^2^) | 94.16 (0.65) | | 92.72 (1.30) | | 95.59 (1.36) | | 0.17 |
| Chronic kidney disease |  | 102 (7.89) |  | 28 (8.67) |  | 24 (7.43) | 0.66 |
| **Polyaromatic hydrocarbons (PAHs)** | **total, n = 1281** | | **1st quartile, n = 320** | | **4th quartile, n = 321** | |  |
| **Characteristic** | **mean (SE)** | **n (%)** | **mean (SE)** | | **mean (SE)** | **n (%)** | **p-value** |
| Age | 47.84 (0.49) | | 50.24 (1.03) | | 45.92 (91.45) | | 0.0018 |
| Male sex |  | 693 (54.10) |  | 167 (52.19) |  | 173 (53.89) | 0.72 |
| Race (non-Hispanic black) |  | 202 (16.70) |  | 32 (10.00) |  | 68 (21.18) | 0.00015 |
| Ratio of family income to poverty level | 2.52 (0.05) | | 2.95 (0.09) | | 1.97 (0.09) | | 6.67E-14 |
| Smoker |  | 336 (26.23) |  | 13 (4.06) |  | 205 (63.86) | 2.20E-16 |
| Weekly alcoholic beverages | 1.29 (0.05) | | 1.21 (0.10) | | 1.37 (0.12) | | 0.3 |
| Body mass index | 28.91 (0.18) | | 27.58 (0.32) | | 28.97 (0.34) | | 0.0033 |
| Diabetes mellitus |  | 137 (10.69) |  | 38 (11.88) |  | 36 (11.21) | 0.89 |
| Systolic blood pressure (mm Hg) | 122.39 (0.51) | | 123.56 (1.06) | | 122.25 (1.04) | | 0.38 |
| Diastolic blood pressure (mm Hg) | 69.61 (0.36) | | 69.43 (0.77) | | 69.44 (0.75) | | 0.99 |
| serum creatinine (mg/dL) | 0.91 (0.01) | | 0.91 (0.01) | | 0.89 (0.02) | | 0.44 |
| estimated glomerular filtration rate (mL/ minute/ 1.73 m^2^) | 94.23 (0.65) | | 90.69 (1.27) | | 97.60 (1.27) | | 0.00013 |
| Chronic kidney disease |  | 101 (8.51) |  | 34 (10.63) |  | 20 (6.23) | 0.063 |
| **High molecular-weight (HMW) phthalates** | **total, n = 1292** | | **1st quartile HMW, n = 323** | | **4th quartile HMW, n = 323** | |  |
| **Characteristic** | **mean (SE)** | **n (%)** | **mean (SE)** | **n (%)** | **mean (SE)** | **n (%)** | **p-value** |
| Age | 47.89 (0.49) | | 50.39 (1.00) | | 45.22 |  | 1.90E-04 |
| Male sex |  | 699 (54.10) |  | 163 (50.46) |  | 189 (58.51) | 0.048 |
| Race (non-Hispanic black) |  | 202 (15.63) |  | 43 (13.31) |  | 57 (17.65) | 0.16 |
| Ratio of family income to poverty level | 2.52 (0.05) | | 2.59 (0.09) | | 2.54 (0.10) | | 0.73 |
| Smoker |  | 335 (25.93) |  | 86 (26.63) |  | 82 (25.39) | 0.79 |
| Weekly alcoholic beverages | 1.29 (0.05) | | 1.41 (0.12) | | 1.17 (0.10) | | 0.14 |
| Body mass index | 28.92 (0.18) | | 27.61 (0.31) | | 29.81 (0.38) | | 7.38E-06 |
| Diabetes mellitus |  | 139 (10.76) |  | 28 (8.67) |  | 40 (12.38) | 0.16 |
| Systolic blood pressure (mm Hg) | 122.43 (0.51) | | 124.83 (1.14) | | 121.16 (0.93) | | 0.01 |
| Diastolic blood pressure (mm Hg) | 69.58 (0.36) | | 69.07 (72) |  | 69.47 (0.77) |  | 0.71 |
| serum creatinine (mg/dL) | 0.91 (0.01) | | 0.92 (0.02) | | 0.91 (0.02) | | 0.86 |
| estimated glomerular filtration rate (mL/ minute/ 1.73 m^2^) | 94.16 (0.65) | | 91.59 (1.32) | | 96.18 (1.29) | | 0.01 |
| Chronic kidney disease |  | 102 (7.89) |  | 30 (9.29) |  | 27 (8.36) | 0.78 |
| **Low molecular-weight (LMW) Phthalates** | **total, n = 1292** | | **1st quartile LMW, n = 323** | | **4th quartile LMW, n = 323** | |  |
| **Characteristic** | **mean (SE)** | **n (%)** | **mean (SE)** | **n (%)** | **mean (SE)** | **n (%)** | **p-value** |
| Age | 47.89 (0.49) | | 47.45 (0.96) | | 47.72 (0.94) |  | 9.40E-01 |
| Male sex |  | 699 (54.10) |  | 177 (54.80) |  | 182 (56.35) | 0.92 |
| Race (non-Hispanic black) |  | 202 (15.63) |  | 15 (4.64) |  | 81 (25.08) | 6.49E-13 |
| Ratio of family income to poverty level | 2.52 (0.05) | | 2.87 (0.10) | | 2.43 (0.09) | | 7.68E-04 |
| Smoker |  | 335 (25.93) |  | 75 (23.22) |  | 29.72 | 0.07 |
| Weekly alcoholic beverages | 1.29 (0.05) | | 1.47 (0.11) | | 1.17 (0.10) | | 0.056 |
| Body mass index | 28.92 (0.18) | | 27.31 (0.31) | | 29.90 (0.38) | | 1.48E-07 |
| Diabetes mellitus |  | 139 (10.76) |  | 28 (8.67) |  | 33 (10.22) | 0.59 |
| Systolic blood pressure (mm Hg) | 122.43 (0.51) | | 121.90 (1.00) | | 122.93 (1.10) | | 0.49 |
| Diastolic blood pressure (mm Hg) | 69.58 (0.36) | | 69.25 (0.71) | | 70.73 (0.79) |  | 0.16 |
| serum creatinine (mg/dL) | 0.91 (0.01) | | 0.95 (0.03) | | 0.89 (0.01) | | 0.085 |
| estimated glomerular filtration rate (mL/ minute/ 1.73 m^2^) | 94.16 (0.65) | | 92.26 (1.33) | | 96.16 (1.24) | | 0.03 |
| Chronic kidney disease |  | 102 (7.89) |  | 29 (8.98) |  | 19 (5.88) | 0.18 |
